# Reconfiguration of Dominant Coupling Modes in Mild Traumatic Brain Injury Mediated by δ-band Activity: a Resting State MEG Study

**DOI:** 10.1101/142117

**Authors:** Marios Antonakakis, Stavros I. Dimitriadis, Michalis Zervakis, Andrew C. Papanicolaou, George Zouridakis

**Author notes:** **Corresponding author:** Marios Antonakakis, PhD Student, Institute for Biomagnetism and Biosignal Analysis, Westfalian Wilhelms-University Muenster, Malmedyweg 15, 48149 Muenster, Germany.

## Abstract

During the last few years, rich-club (RC) organization has been studied as a possible brain-connectivity organization model for large-scale brain networks. At the same time, empirical and simulated data of neurophysiological models have demonstrated the significant role of intra-frequency and inter-frequency coupling among distinct brain areas. The current study investigates further the importance of these couplings using recordings of resting-state magnetoencephalographic activity obtained from 30 mild traumatic brain injury (mTBI) subjects and 50 healthy controls. Intra-frequency and inter-frequency coupling modes are incorporated in a single graph to detect group differences within individual rich-club subnetworks (type I networks) and networks connecting RC nodes with the rest of the nodes (type II networks). Our results show a higher probability of inter-frequency coupling for (δ−γ_1_), (δ−γ_2_), (θ−β), (θ−γ_2_), (α−γ_2_), (γ_1_−γ_2_) and intra-frequency coupling for (γ_1_−γ_1_) and (δ−δ) for both type I and type II networks in the mTBI group. Additionally, mTBI and control subjects can be correctly classified with high accuracy (98.6%), whereas a general linear regression model can effectively predict the subject group using the ratio of type I and type II coupling in the (δ, θ), (δ, β), (δ, γ_1_), and (δ, γ_2_) frequency pairs. These findings support the presence of an RC organization simultaneously with dominant frequency interactions within a single functional graph. Our results demonstrate a hyperactivation of intrinsic RC networks in mTBI subjects compared to controls, which can be seen as a plausible compensatory mechanism for alternative frequency-dependent routes of information flow in mTBI subjects.

## Introduction

While traumatic brain injury (TBI) is one of the most serious brain disorders, mild TBI (mTBI) is one of the most frequent, and accounts for almost 90% of all brain injuries (Levin et al., 1987; Len and Neary, 2011; Huang et al., 2014). The symptomatology of brain injury is characterized by headaches, fatigue, memory loss, sleep disturbances, loss of balance, seizures, depression, and visual and emotional disturbances (Huang 2014). It is estimated that 5-20 percent of irremediable patients (Bharath et al., 2015) have symptoms that persist for one year or more after the injury (Huang et al. 2014). Based on these findings, a number of research groups have worked on developing robust biomarkers for highly accurate differentiation of mTBI patients from healthy controls using resting state magnetoencephalographic (MEG) recordings and functional brain connectivity analysis (Huang et al., 2009, 2014; Zouridakis et al., 2012; Antonakakis et al., 2016a; Dimitriadis et al., 2015a, Mvula et al., 2017; Zouridakis et al., 2017).

In terms of brain communication, both structural and functional imaging studies have shown (van den Heuvel and Sporns, 2011, Palva and Palva, 2011; Vértes and Bellmore, 2015) that the highest amount of information flows within a backbone of the brain network consisting of a subset of main nodes, or hubs, known as rich club (RC) that often follows a small-world (SW) topology. The SW network model has been investigated in Alzheimer’s disease (Stam et al., 2007), schizophrenia (Micheloyannis et al., 2006), and autism (Liu et al., 2008; Rubinov et al., 2010; Tsiaras et al., 2011), whereas the RC organization has been observed both in computer simulations (Senden et al., 2014) and human studies involving healthy subjects (van den Heuvel and Sporns, 2011; Bullmore and Sporns, 2012; Mišić et al., 2014), as well as brain ischemia (Fornito et al., 2012; van den Heuvel and Sporns, 2013; Crossley, Alawieh, Watanabe et al., 2015) and mTBI patients (Antonakakis et al., 2015). RC nodes play a significant role in communication and information integration among brain areas that are distinct and distant. Thus, it is important to explore how this integration of information is affected by various brain diseases and disorders (van den Heuvel and Sporns, 2011; Mišić et al., 2014; Bullmore and Sporns, 2012).

Functionally, the human brain consists of several specialized subsystems, each oscillating in a dominant frequency. Communication between a small and a larger system is facilitated via intra-frequency coupling, whereas communication between two larger systems, whereby each system oscillates with its own prominent frequency, is realized via cross-frequency coupling (Canolty et al., 2006). A key feature of ongoing brain activity is its intrinsic coupling mode which exhibits multiple spatio-temporal patterns and supports rich information processing (Varela et al., 2001). There is significant evidence that these intrinsic coupling modes are negatively affected by brain diseases and positively reinforced by cognition and learning (Engel et al., 2013).

Important issues stemming from previous analyses on the study of mTBI using Granger causality (Zouridakis et al., 2012), phase synchronization (Dimitriadis et al., 2015b), cross-frequency coupling (Antonakakis et al., 2015, 2016a, c), complexity (Antonakakis et al., 2016b), as well as brain activation patterns of both EEG and MEG at the sensor (Li et al., 2015) and source (Zouridakis et al., 2016; Li et al., 2017) levels relate to the following key questions: 1) Is there a group difference in intra-frequency and inter-frequency coupling within the RC networks (type I network) and between the RC hubs and the rest of the brain network (type II network)? 2) If so, in which intra-frequency and inter-frequency intrinsic coupling modes does the ratio of probability distributions between the two types of networks show group differences? 3) Are the theoretical information exchange rate (IER), the weighted IER (WIER), and the ratio of probabilities between the two types of networks altered in mTBI? 4) Can the ratio of probability distribution of the prominent intrinsic coupling modes between the two types of networks discriminate the two groups? To address these questions, in the current study we explore both intra-frequency and inter-frequency coupling using resting-state MEG obtained from mTBI patients and healthy controls under the distinction of brain network nodes as RC and non-RC hubs.

The present study is structured as follows: the next section describes the Experimental Procedures including the subjects and analysis methods, the subsequent section presents the analysis results, whereas the last section discusses advantages and limitations of the proposed methodology and describes further future improvements.

## Experimental Procedures

### Participants and procedure

Thirty right-handed mTBI patients (29.33 ± 9.2 years of age) (Levin, 2009) and fifty age- and gender-matched neurologically intact healthy controls (29.25 ± 9.1 years of age) participated in the study. The control group was drawn from a normative data repository at UTHSC-Houston, whereas the mTBI patients were recruited from three trauma centers in the greater Houston metropolitan area. Those centers were part of a larger study (Levin, 2009) supported by Department of Defense (DoD). mTBI was defined according to the guidelines of DoD (Assistant Secretary, 2007) and the American Congress of Rehabilitation Medicine (Kay et al., 1993). Demographic details about the mTBI patients are presented in the Supplementary Material, which includes all information provided by the clinicians. Previous head injuries, history of neurologic or psychiatric disorder, substance abuse, and extensive dental work and implants incompatible with MEG were used as exclusion criteria for the control group. The project was approved by the Institutional Review Boards (IRBs) at the participating institutions and the Human Research Protection Official’s review of research protocols for DoD. All procedures were compliant with the Health Insurance Portability and Accountability Act (HIPAA).

The MEG acquisition included ten minutes of resting-state activity for each subject lying on a bed with eyes closed, using a whole-head Magnes WH3600 system with 248 channels (4D Neuroimaging Inc., San Diego, CA). Data were acquired using a sampling rate of 1017.25 Hz and online bandpass filters between 0.1–200 Hz. Five minutes of data were artifact contaminated (Dimitriadis et al. 2015a) and thus the rest five minutes were used in the current analysis. The original axial gradiometer recordings were transformed to planar gradiometer field approximations using the *sincos* method implemented in the software package Fieldtrip (Oostenveld et al., 2011).

### MEG Preprocessing

Reduction of non-cerebral activity was based on an automated blind detection and elimination strategy applied to the raw MEG data, due to the lack of independent ocular and cardiac activity monitoring, using the Fieldtrip toolbox (Oostenveld et al., 2011) and MATLAB (The MathWorks, Inc., Natick, MA, USA). In particular, the following iterative procedure was applied to all datasets individually: First, correction of activity from bad MEG channels was performed using interpolation (Oostenveld et al., 2011) on the four closest channels surrounding the bad one, whereas notch filtering was used to eliminate the effects of power line noise at 60 Hz. Second, blind detection of non-cerebral activity relied on Independent Component Analysis or ICA (Delorme and Makeig, 2004) and information theory metrics. Detection started using principal component (PC) analysis to eliminate external magnetic noise, whitening of brain activity, and reducing the dimensionality of the original data using principal component analysis. A threshold of 95% of the total variance (Delorme and Makeig, 2004; Escudero et al., 2011; Antonakakis et al., 2015, 2016a) was used to select the optimum number of PCs. Then, the reduced number of PCs were fed to the Infomax algorithm (Delorme and Makeig, 2004) to identify the independent components (ICs). Subsequently, elimination of IC corresponding to artifactual activity was done using kurtosis, Rényi entropy, and skewness on the entire time course of each IC, separately for each subject. These values were normalized to zero-mean and unit-variance. Then, an IC was tagged as representing ocular or cardiac activity if more than 20% of kurtosis, Rényi entropy, and skewness were outside the range [−2, +2] (Escudero et al., 2011; Dimitriadis et al., 2013a; Antonakakis et al., 2015, 2016a). Additionally, we used the time course of each IC, its spectrum profile, and the topological distribution of the IC weights to further confirm if an IC was an artifact. Across subjects, the number of ICs removed was on average 6 out of 50 ICs. The artifact-free ICs were then used to reconstruct the MEG activity.

### Functional Connectivity Graphs

To estimate intra-frequency and inter-frequency connections, the artifact-free multidimensional array **X** (sensors × time series) was filtered in six standard brain frequency rhythms/bands. In particular, for each subject, **X** was bandpass filtered in the δ (0.5 − 4 Hz), θ (4 − 8 Hz), α (8 − 15 Hz), β (15 − 30 Hz), γl (30 − 45 Hz), and γ2 (45 − 80 Hz) using a fourth order two-pass Butterworth filter. This resulted in a multidimensional array, **X**_f_, where f = δ, θ, α, β, γ_1_, and γ_2_.

Intra-frequency functional connectivity graphs (IFCG) were constructed using the non-linear metric mutual information (MI), which expressed the intra-frequency content between MEG time-series in a brain rhythm. In addition, cross-frequency interactions were explored by analyzing inter- or cross-frequency functional connectivity graphs (CFCG) based on phase-to-amplitude coupling (PAC) for the inter-frequency content within a single MEG time series or between pairs of MEG time series. As a result of the IFCG and CFCG, the so-called *ICFCG* was derived via surrogate analysis such as to reveal dominant intra-frequency or inter-frequency coupling modes for each pair of MEG sensors. The rich club organization showed statistically significant differences and separation of the two group based on the theoretical amount of information and probability density functions.

### IFCG – Mutual Information

IFCG were constructed using MI revealing the interdependence between MEG time series X_f,i_ and X_f,j_ (i,j = 1 … 248) of **X**_f_. The use of MI stems from information theory and offers several advantages, such as sensitivity to any type of dependence between the time series, including nonlinear relations and generalized synchronization, robustness to outliers, and measurement in bits. The mathematical definition of MI between two specific-band, artifact-free, filtered sensor data arrays X_f,i_ and X_f,j_ is given by

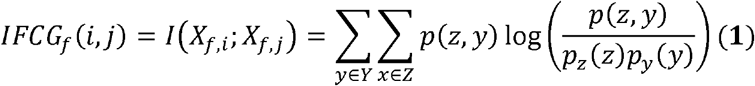

where Z = X_f,i_, Y = X_f,j_, p(z,y) is the joint probability distribution function of Z and Y, respectively, and p_z_(z) = Σ_y∈Y_P(z,y) and P_y_(y) = Σ_z∈Z_P(z,y) are the marginal probability distribution functions of Z and Y, respectively (Tsiaras et al., 2011; Antonakakis et al., 2015).

### CFCG — Cross-frequency Coupling

In CFCG, PAC was used to reveal the relation of low- and high-pass frequency content within an MEG sensor or between pairs of MEG sensors for f_c_= (δ, θ), …, (γ_1_, γ_2_). In terms of PAC, the phase of low-frequency rhythm modulated the amplitude of a higher-frequency oscillation (Tort et al., 2008; Voytek et al., 2010; Xu et al., 2013). PAC was calculated between MEG sensors X_i_ and X_j_ (i, j = 1 … 248) of a multidimensional array of time series **X** using MI. Specifically, Eq. (1) was used between the phases of low-frequency (f_i_) versions of the MEG sensors. First, the phase ϕf_l,i_ of the low-frequency content of X_i_, extracted by Hilbert transform, was used as *Z* = *φ_fl,l_*,. The corresponding low-pass phase, computed by the amplitude X_fh,j_ of the high-frequency (f_h_) content of X_j_, was used as *Y* = *φ_fh,fl,j_*. More details are completely described in Supplementary Material.

### ICFCG estimations

The based FCG type, *ICFCG (combination of IFCG and CFCG)*, was estimated using intra-frequency and inter-frequency couplings. The ICFCG contains the dominant frequency mode (intra- or inter-) of each MEG sensor. In particular, the mathematical definition of this type of FCG is given by

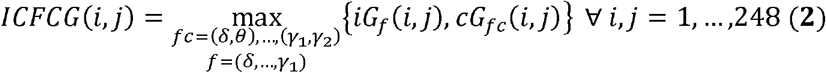

where *iG_f_* is the IFCG and *cG_fc_*(*i*,*j*) is the CFCG including all frequency pairs (i.e., 15 frequency pairs) and all frequencies (i.e., 6 frequencies) for each pair of MEG sensors.

### Surrogate analysis

A surrogate data analyses was employed to identify significant intra- and inter-frequency interactions which were estimated for every frequency and pair of frequencies, respectively, within and between the 248 sensors (Theiler et al., 1992). Thus, it was possible to determine (a) if a given MI value differed from what would be expected by chance alone, and (b) if a given non-zero MI value indicated synchronization that was, at least statistically, non-spurious.

The null hypothesis H_0_ stated that the observed MI value came from the same distribution as the distribution of surrogate MI values for every sensor pair, frequency, and frequency pair independently. One thousand surrogate time-series *MI*(*t*) were generated by cutting at a single point at a random location and exchanging the two resulting time courses (Canolty et al., 2006; Aru et al., 2015). Repeating this procedure produced a set of surrogates with minimal distortion of the original synchronization dynamics and impact on the non-stationarity of brain activity as compared to either merely shuffling the time series or cutting and rebuilding the time series in more than one time points. This procedure ensures that the observed and surrogate indices shared the same statistical properties. For each data set, the surrogate MI (S_MI_) was computed. We then determined a one-sided p-value expressing the likelihood that the observed MI value could belong to the surrogate distribution, and corresponded to the proportion of “surrogate” Mis which was higher than the observed MI value (Theiler et al., 1992). MI values associated with statistically significant p-values were considered unlikely to reflect signals not entailing MI coupling.

A similar procedure was adopted for CFCG and ICFCG. Regarding the ICFCG, we define the dominant type of interaction in intra-frequency and inter-frequency coupling by first correcting the p-value level (p) using Bonferroni correction (IFCG: p’ = p/(6 frequency bands), CFCG: p’ = p/(15 frequency pairs) and ICFCG: p’ = p/(21=(6 (intra-frequency) + 15 (inter-frequency couplings)). This statistical thresholding scheme could result in three possible outcomes:

1. only one p’-value exceeded the threshold, in which case we assigned the related coupling mode (intra-frequency e.g., delta, or inter-frequency: e.g., delta-theta) to this pair of MEG sensors;
2. more than two p’-values exceeded the correction, in which case we assigned the one with maximum MI value to this pair of MEG sensors; or
3. none p’-value crossed the threshold, in which case we assigned zero to the particular pair of MEG sensors.

Then, the false discovery rate (FDR) adjustment (Benjamini and Hochberg, 1995) was employed to control for multiple comparisons across all combinations of sensor pairs, independently for each frequency and frequency pair, with the expected proportion of false positives set to q ≤ 0.01. Finally, only the significant connections were kept with their MI weights while the rest were substituted with zeros.

### Topological Filtering

Each of the brain connectivity graphs described — IFCG, CFCG, and ICFCG — resulted in a k x k matrix of connectivity values (k is the number of the MEG sensors) representing a fully connected, weighted, symmetric, directed FCG. To reduce the maximum number of possible connections in the FCGs (k=248 leads to k^2^= 61504 possible connections) and allow only patterns with the most topologically significant connections to emerge, the actual connections were filtered using a data-driven topological thresholding scheme based on global information among the sensor links (Bassett et al., 2009; Dimitriadis et al., 2015a). We applied this approach on each type of FCG for all subjects. The filtering procedure is described elsewhere (Dimitriadis et al., 2015a, Antonakakis et al., 2016a) and the corresponding software implementation is available online^1^.

### Rich Club Estimation

The RC organization was estimated for all FCG types using the Brain Connectivity Toolbox^2^ (Rubinov and Sporns, 2010). One thousand random networks preserving the degree distribution and sequence of the original network (van den Heuvel and Sporns, 2011) were generated and the RC coefficient was computed for each random network and degree k. Φ_r_^w^ was computed as the average RC coefficient over the random networks and the normalized RC parameter Φ_n_^w^ was computed as the ratio of Φ^w^to Φ_r_^w^. The randomization process could be used to assess the statistical significance of the results through permutation testing (van den Heuvel and Sporns, 2011). In that respect, the distribution of Φ_r_^w^ yielded the null distribution of RC coefficients obtained from random topologies. Using this null distribution, Φ^w^ could be assigned a p-value from the percentage of random tests found to be more extreme than the observed RC coefficient Φ^w^. All tests were performed at the FDR-adjusted p’ level of significance (Benjamíni and Hochberg, 1995) computed as p’ = p/(80 subjects × 6 frequency bands) for the IFCGs, p’ = p/(80 subjects × 15 frequency pairs) for CFCGs, and p’ = p/(80 subjects) for CFCGs.

### Comodulograms

Comodulograms are matrices tabulating the probability distribution (PD-comodulogram) of connections within a functional connectivity network associated with intra-frequency coupling (diagonal) and inter-frequency coupling (upper diagonal). To estimate the prominent type of interaction for each pair of sensors, across six (intra-) + 15 (inter-) = 21 MI coupling strengths, a surrogate analysis was followed. PD-comodulograms were computed in the range between 0.5 and 80Hz, within and across the eight frequency bands studied, separately for each subject group and type of RC network, as described in the following sections.

### Information Exchange Rate (IER)

A novel measure to summarize the rate of information transfer among neural assemblies throughout the brain was adopted under the assumption that phase-to-amplitude coupling, or PAC, modes reflect processes for exchanging “packets of neural information” among populations of neurons operating at different characteristic frequencies. The concept behind PAC interactions can be interpreted as the process of forming “packets of information,” in which higher frequency brainwaves are nested within the phase of slower rhythms. A specific number of cycles of the higher frequency oscillation can be incorporated within the phase of the slower frequency. This number is expected to reflect the amount of information that can be exchanged among neural oscillators operating at different characteristic frequencies. With this assumption, and based on the detected prominent cross-frequency interactions, we adopted a previously introduced measure that aggregates the rate of information exchanged throughout the brain (Dimitriadis et al., 2016b): for each subject, we simply summed up the number of cycles of the higher frequency that could be included within the phase of the slower frequency. This index, which provides the “instantaneous” information exchange rate (IER), varies between 0 and 1 and is defined as follows,

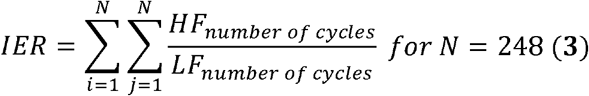

Since each of the detected PAC interactions is associated with a varying strength, or a MI level, we also introduced a “weighted” version of the above index, which also ranges between 0 and 1 and is defined as follows (Dimitriadis et al., 2016b),

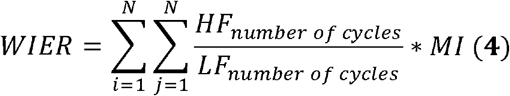

The WIER magnitude can be interpreted as the co-modulations between lower and higher frequencies, with 1 reflecting the strongest PAC interaction. PAC value can be used as an indicator of how active a “channel” is between or within sensors for information exchange.

### Network Features Related to the Dominant Type of Interaction

Many possible network features can be extracted from brain networks based on different types of interaction in the RC organization. We estimated the distribution of RC hubs over several brain areas in both subject groups, and connectivity graph types, i.e., IFCG, CFCG, and ICFCG. Furthermore, by dividing the entire network into two subnetworks, i.e., the rich-club subnetwork and the one composed of connections between RC nodes and the rest of the network, different features can be evaluated. Namely, the first or type I network is that with connections among RC nodes, while the second or type II network, is the one connecting the RC nodes to the rest of the network nodes. Afterwards, we estimated the ratio of type I network PD-comodulogram divided by the type II network PD-comodulogram based on ICFCGs.

### Exploration of Statistical Differences

Statistical analysis was performed on the IER and wIER values as well as on the corresponding ratio of type I to type II network and on the ratio of type I to type II network PD-comodulogram to detect possible significant differences between the two groups. The statistical methods used included normality test and parametric and non-parametric pairwise tests and were similar to our previous study (Antonakakis et al., 2016a). The threshold for significance of the p-value was set to 95%. After FDR adjustment (Benjamini and Hochberg, 1995) the new p’ values where given by p’ = p/2 for IER, wIER, and their corresponding ratio of type I and II networks, and p’ = p/21 (accounting for six frequency bands and 15 frequency pairs).

## Results

### Probability Distribution of RC Hubs over Brain Regions

In an attempt to consistently estimate the spatial distribution of RC hubs over each group and FCG type (IFCG, CFCG, and ICFCG), we integrated their representation over distinct brain regions in both hemispheres (frontal, central, temporal, parietal, and occipital). In particular, we measured the discrete probabilities for RC hubs across regions, separately for each subject, as the ratio of the number of RC nodes in a specific brain area to the total number of RC nodes detected for that subject.

The corresponding averaged distributions are depicted across group and FCG type in Fig. 1. Regarding the IFCG-RC topology, RC hubs with higher probability density were detected in both groups over frontal and temporal regions bilaterally in all frequency bands (Fig. 1a) and with lower probability density value in the other regions. Significant differences (p<0.05, Bonferroni corrected, p’<p/48 (6 frequency bands x 8 lobes)) were found in left tempo-parietal regions in the δ and β frequency bands, and in the right frontal regions in the θ, α, β, and γ_1_ bands. Similar distributions were found for CFCG-RC and IFCG-RC topologies for both groups as seen in Fig. 1b (p<0.05, Bonferroni corrected, p’<p/120 (15 frequency bands x 8 regions)). Most of the significant differences were seen in parietooccipital regions in all frequency pairs. Finally, even though the ICFCG-RC of Fig. 1C looked similar to IFCG-RC, no significant differences were found.

**Figure 1.**
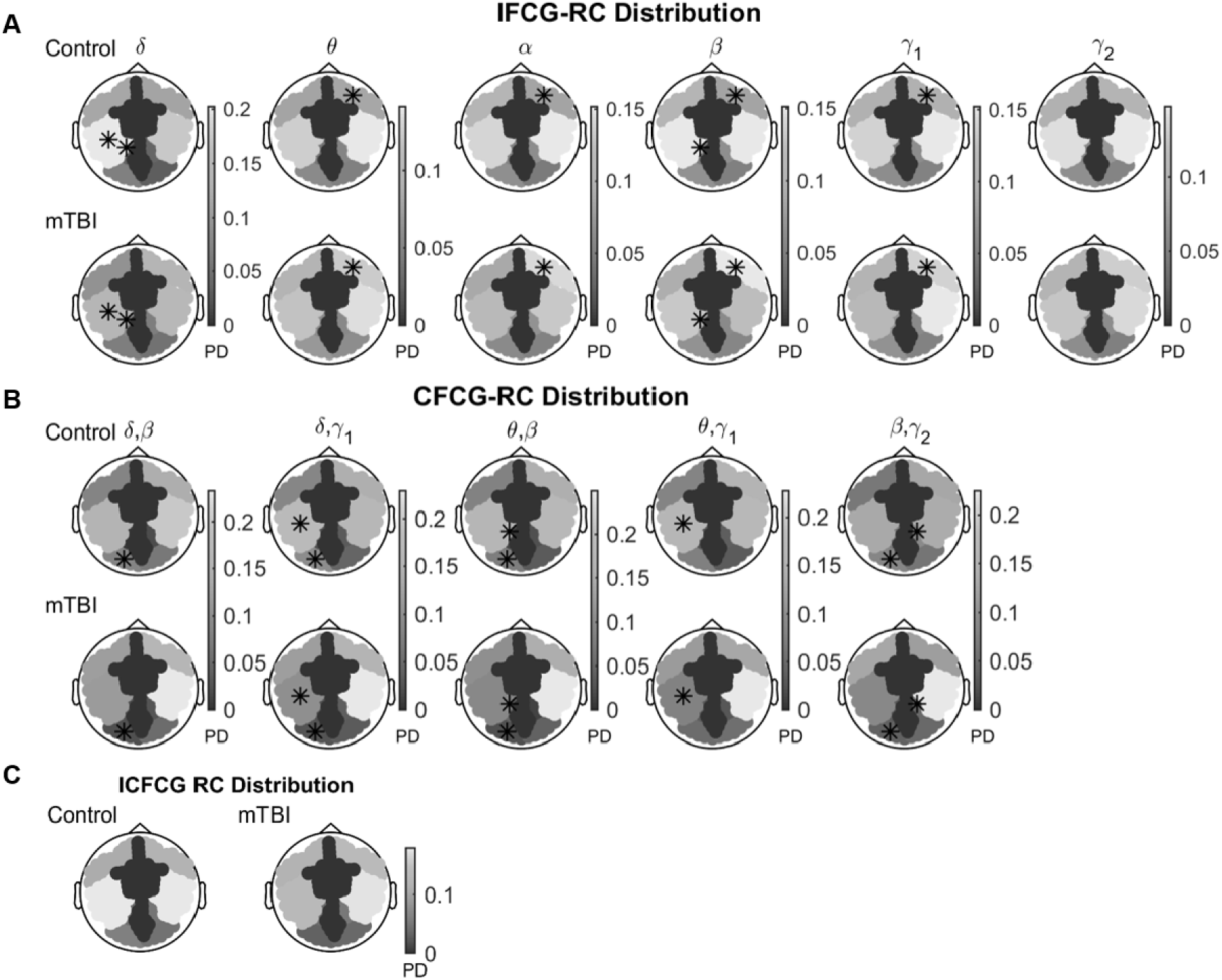
Average distribution of RC nodes in mTBI and Control subjects for different types of functional connectivity graphs: **A**) intra-connections (MI – IFCG), **B**) inter-connections (CFC – CFCG), and **C**) intra – CFC connections (ICFCG). In the case of **A**) five inter-frequency pairs show classification accuracy higher than 90%. The colorbar common for both groups denotes propability density. A black * indicates statistically significant differences (p’<0.05) between the two groups over a particular brain region.

### Comodulograms of Dominant Intrinsic Coupling Modes based on RC Subnetwork

Figure 2 shows the PD-comodulogram matrices for connections within networks associated with intra-frequency (diagonal) and inter-frequency coupling (upper diagonal). The ratios of the type I to type II network comodulograms are shown in Fig. 2a and Fig. 2b, respectively, for the two groups. Namely, significant differences (p < 0.01, (p<0.05, Bonferroni corrected, p’<p/21 – black ‘**’) were observed in the δ, α, and γ_1_ low frequency modulating phase and δ and γ_2_ high frequency amplitude. Less significant differences (p < 0.05, Bonferroni corrected, p’<p/21 – black ‘*’) were observed for δ, θ, and γ_1_ low frequency modulating phase and γ_1_, β, and γ_1_ high frequency amplitude.

**Figure 2.**
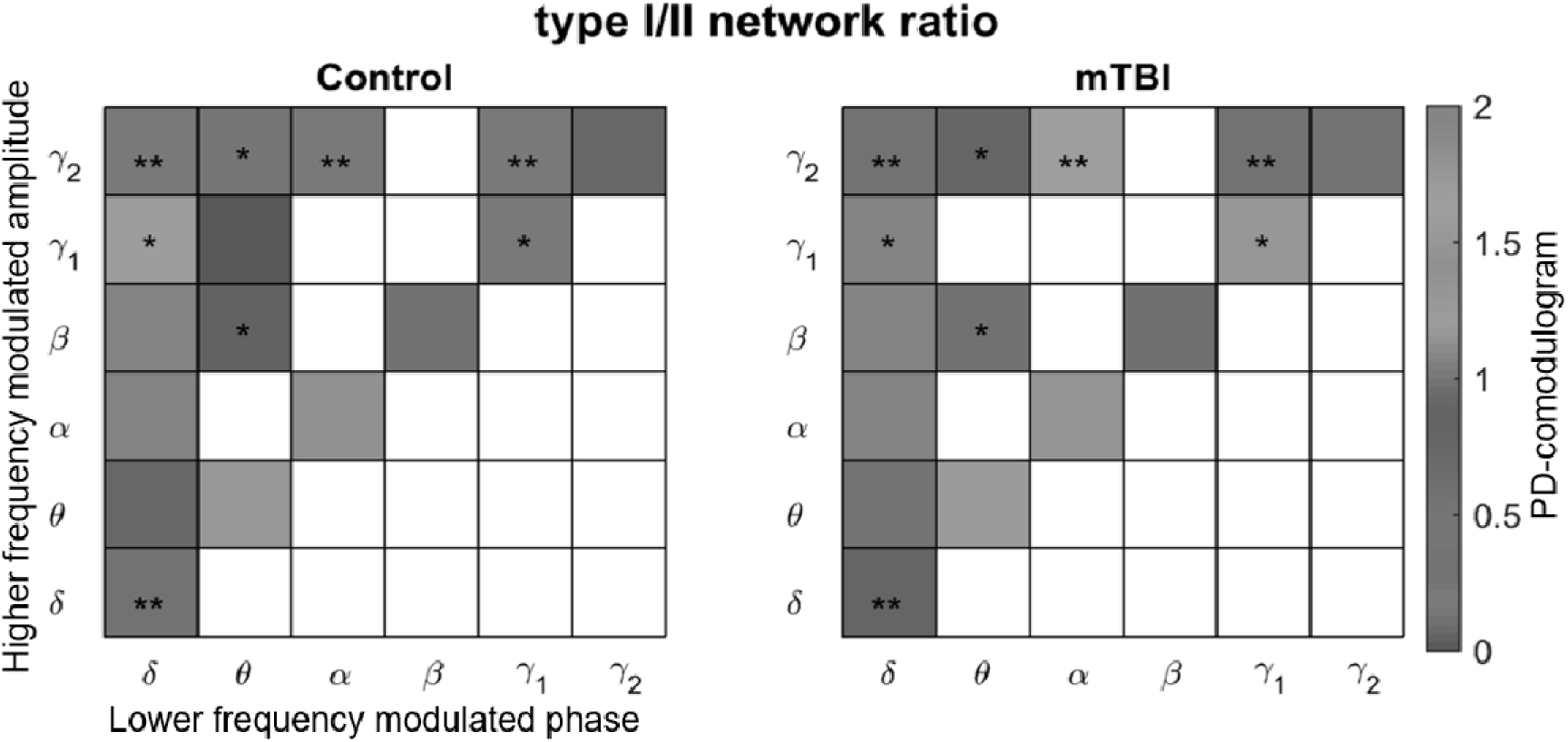
Ratio of comodulogram of type I to type II network for control (left) and mTBI (right) subjects. The horizontal axis encodes the modulating phase of the lower frequency and the vertical axis reflects the modulated amplitude of the higher frequency. Different colors encode CFCG strength between frequency pairs; black‘*’ and ‘**’ denote statistical significance levels of p’< 0.05 and p’< 0.01, respectively.

### Theoretical IER/WIER based on RC Subnetwork

We defined the information exchange ratio (IER) as the sum of ratios between the amplitude of f_H_ and the phase of f_L_ to quantify the theoretical amount of information exchanged among RC hubs according to the dominant coupling mode. Figure 3 presents the averaged values across each group of IER, WIER and their corresponding ratio of type I to type II networks (IER_ratio_ and WIER_ratio_). The control group showed higher IER (Fig. 3a) and statistically significant WIER values (Fig. 3b) than the mTBI group. However, the IER_ratio_and WIERrat¡o metrics (Fig. 3c) were higher in the mTBI group compared to the control group.

**Figure 3.**
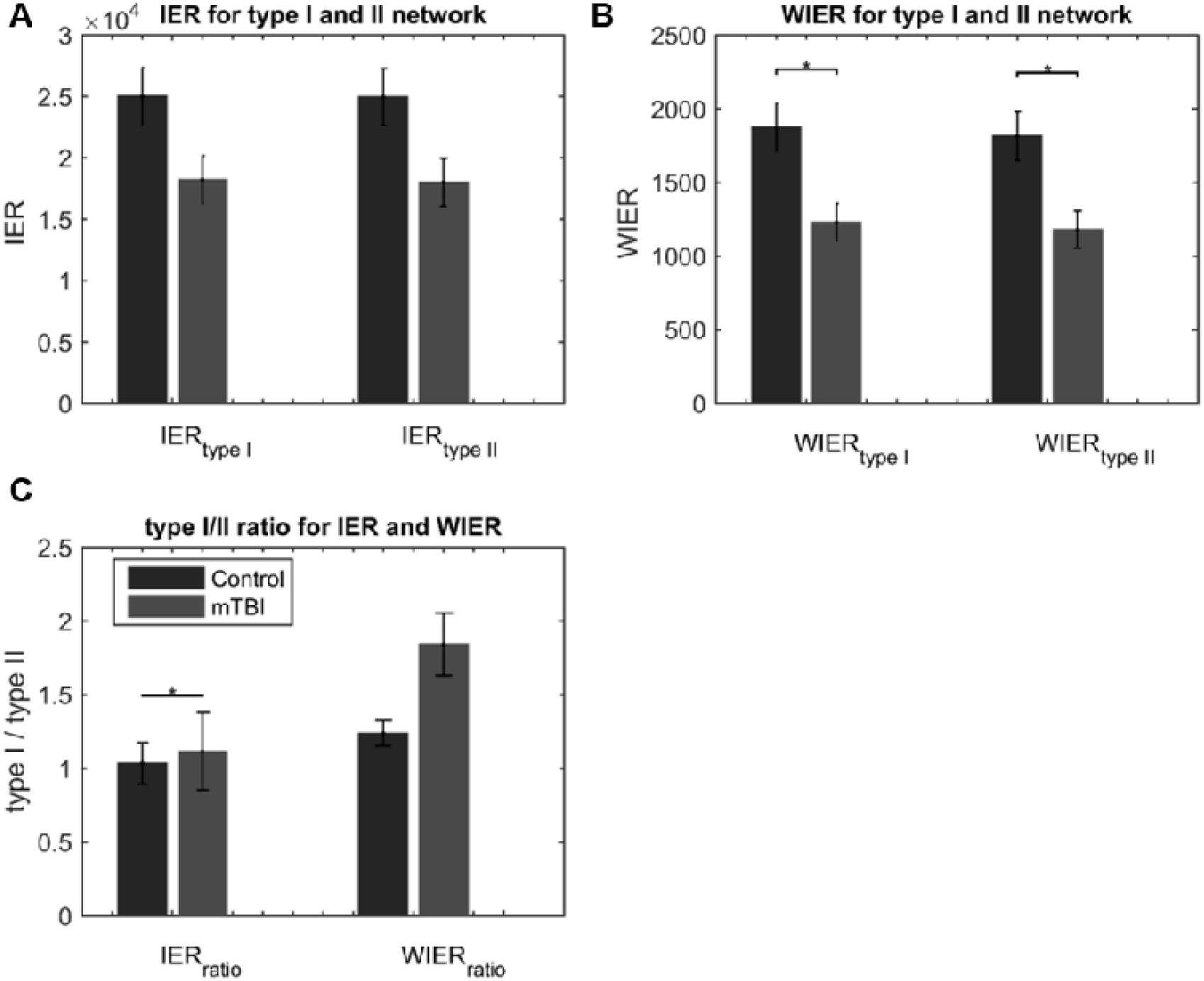
Theoritical amount of information **A**) IER and **B**) WIER for type I and type II networks. **C**) The averaged ratio of type I/type II for IER and WIER for mTBI and Control subjects. All comparisons (paired test linked by *) reach statistical significance (p’: * < 0.05; ** < 0.01 and ***<0.001).

Overall, we found a hyperactivity within RC subnetwork (type I network) for mTBI subjects compared to controls, both in the IER_ratio_ and WIER_ratio_ metrics. This hyperactivity was seen in the δ frequency band that modulated γ_1_ and γ_2_, and in the δ, θ, α_1_ and γ_1_ that modulated the γ_2_ band.

### Feature Extraction and Classification Performance

In the current study, we tested the separation of the mTBI and control groups using only ICFCG-based features composed by PD-comodulogram matrices, IER/WIER values, and their corresponding IER_ratio_ and wlER_ratio_. Laplacian scores (LS, He et al., 2005) were estimated through an iterative bootstrap procedure for estimating the cut-off threshold of the features (Dimitriadis et al., 2015a; Antonakakis et al., 2016a, b). Classification evaluation was followed by a 10-fold cross-validation evaluation of one hundred iterations. Two classifiers were used, k nearest neighbor (kNN; Horn and Mathias, 1990) and support vector machine (SVM, Cortes and Vapnik, 1995) to observe the stability of the results.

Table 1 shows that the only surviving features following bootstrap thresholding were the PD-comodulogram values for frequency pairs (δ, θ), (δ, β), (δ, γ_1_), and (δ, γ_2_).

**Table 1.** Classification features after bootstrap thresholding: only the PD-comodulogram values for frequency pairs (δ, θ), (δ, β), (δ, γ_1_), and (δ, γ_2_) survived; a dash “−” is used for features that did not survive.

| Group \ Feature Type | PD-Comodulogram | IER | wIER | $IER_{ratio}$ | $wIER_{ratio}$ |
| --- | --- | --- | --- | --- | --- |
| Controls | $0.00025 \pm 0.0022$ | - | - | - | - |
| mTBIs | $0.0001 \pm 0.00091$ | - | - | - | - |

Table 2 shows the classifier performance in discriminating the mTBI subjects from controls using the kNN and SVM classifiers and 10-fold cross-validation repeated 100 times. Ninety percent of the data were used for training and 10% for testing. Positive labels correspond to the control group and negative labels to the mTBI group. Higher classification accuracy (98.6%) was achieved by the SVM classifier compared to kNN (96.1%). Sensitivity and specificity values were also higher for the SVM than the kNN algorithm. In general, the SVM classifier reached higher performance values, but both were quite efficient in predicting the classification group.

**Table 2:** Classification performance of the k-nearest neighbor (kNN) and support vector machine (SVM) classifiers in separating mTBI patients from controls, based on 100 runs of a 10-fold cross-validation procedure. Ninety percent of the data were used for training and ten percent for testing.

|  | Accuracy (%) | Sensitivity (%) | Specificity (%) |
| --- | --- | --- | --- |
| kNN | 96.1±0.5 | 96±0.006 | 96.3±1.5 |
| SVM | 98.6±0.5 | 98±0.001 | 99.6±1.4 |

To further visualize the separation of the two groups, a 3D visual representation was attempted using the Euclidian distance of the selected features among subjects (Fig. 4a). Then, multidimensional scaling was used to project the multidimensional feature space to 3D, and the convex hull of the resulting ICFCGs was estimated to better visualize the separation of subjects. In general, controls showed higher distance values than mTBI patients, and after 3D projection, mTBI patients showed a larger volume (larger variance) than controls (Fig. 4b).

**Figure 4.**
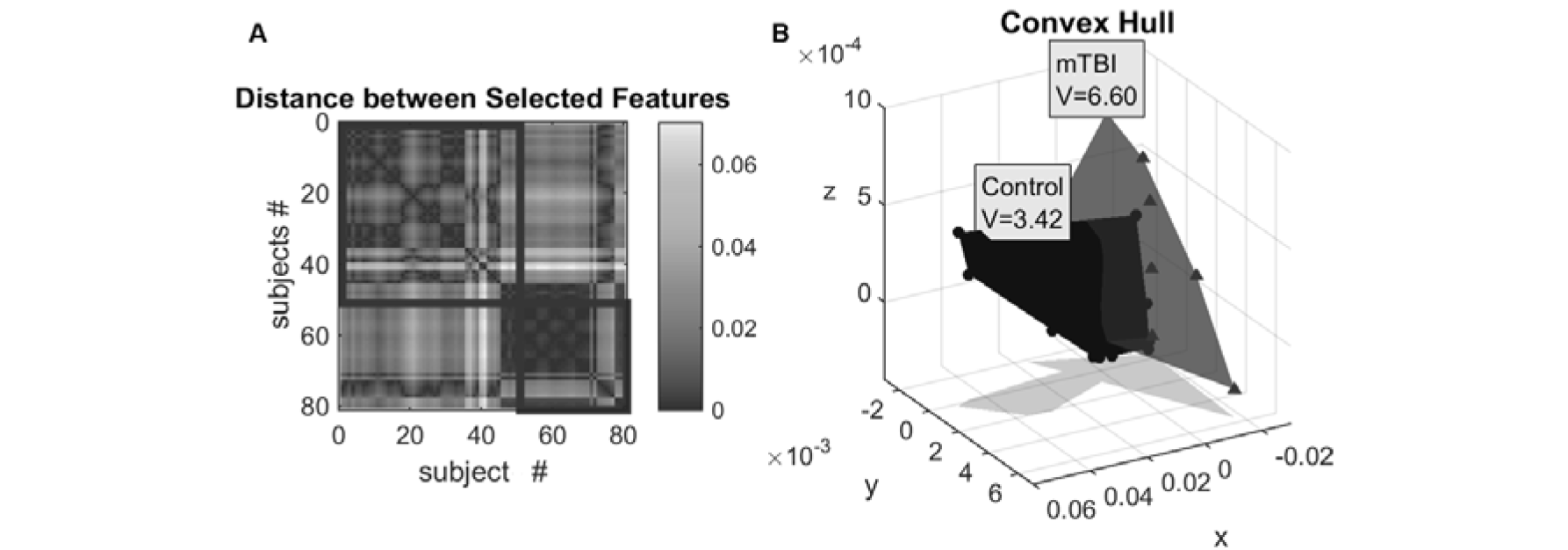
Euclidean distance between classification features. **B**) Convex hull to visualize the separation of the mTBI and Control groups, following multidimensional scaling and 3D projection of the ICFCG selected features (co-modulograms, IER, WIER, IER_ratio_, and WIER_ratio_). Label V denotes the convex hull volume for each group.

A final validation step regarding the significance of the selected features was performed using logistic regression to investigate the group sensitivity with respect to the selected features. Using a general linear regression model of binomial distribution, we tested the linear equation of [*logit*(*Group*) ≈ 1 + (*δ, θ*) + (*δ,β*) + (*δ, γ*_1_) + (*δ,γ*_2_)], where the dependent variable was Group (0,1) and the independent variables were the PD-comodulogram values for frequency pairs (δ, θ), (δ,β), (δ,γ_1_), and (δ,γ_2_) shown in Table 1. Table 3 summarizes the logistic regression results showing that the independent variables are the most significant for the predicting the group. The total number of observations was 80 (50 control and 30 mTBI subjects) with 75 error degrees of freedom, whereas the deviance of fit had 66.8 degrees of freedom with a statistically significant p-value < 0.001. Furthermore, p-values were statistically significant (p< 0.05) for all coefficient estimates, B; thus, all coefficients were informative and could not be rejected. Furthermore, the quite low p-value of the statistical comparison on the model (p-value = 1.09·10^−13^) indicates that this model differs significantly from the constant model (*logit*(*Group*) ≈ 1). The current logistic regression model strongly validates the results of the bootstrap approach regarding the selected features.

**Table 3.** Logistic regression modeling; dependent variable: Group (Control: 0, mTBI: 1); independent variables: PD-comodulogram frequency pairs (δ, θ), (δ, β), (δ, γ_1_), and (δ, γ_2_); B: coefficient estimates; SE: standard error of B; t: *t*-statistic; and p: *p*-values of B.

| Variable | B | SE | t | p |
| --- | --- | --- | --- | --- |
| constant | 169.64 | 76.19 | 22.26 | 0.026 |
| $(\delta, \theta)$ | 6942.1 | 2278.9 | 30.46 | 0.0023 |
| $(\delta, \beta)$ | 1163.7 | 537.03 | 2.167 | 0.031 |
| $(\delta, \gamma_1)$ | -338.53 | 112.35 | -30.13 | 0.0026 |
| $(\delta, \gamma_2)$ | -174.36 | 77.056 | -22.63 | 0.024 |

## Discussion

From the machine learning perspective, we observed a hyperactivity of the type I network compared to type II network in mTBI subjects. The corresponding levels for the control group are shown in Fig.3c, for the IER_ratio_and WIER_ratio_ metrics. The hyperactivity is limited to the δ frequency band, which modulates θ, β, γ_1_, and γ_2_ frequencies (Table 2). The proposed strategy of defining type I and II networks and the subsequent study of prominent intrinsic coupling modes using intra-frequency and inter-frequency estimates succeeded to uncover a hyperactivity for mTBI subjects within the RC module. This hyperactivity can be viewed as a compensatory mechanism that preserves information flow under network disruptions resulting from mTBI. Future follow-up studies should further validate whether the proposed exploratory analysis could be useful for recovery mechanisms (Tarapore et al., 2013) at the individual level of a mTBI patient.

Our analysis introduced several innovative features that succeeded to not only differentiate the mTBI group from controls, but also explain their difference based on network analysis, using appropriate connectivity estimators for both intra-frequency and inter-frequency intrinsic coupling modes. Notice that the two groups are matched in terms of age, but there might be other mismatch between the patient and control groups. The whole analysis procedure is summarized in the following steps:

- Estimate functional brain networks within and between frequency pairs
- Detect the dominant type of interaction for each pair of sensors
- Estimate RC hubs based on the mixed functional connectivity graph for each subject
- Define two subnetworks, within the RC hubs (type I) and between RC hubs and the
rest of the network (type II)
- Estimate the ratio of the PD-comodulograms from dominant types of interactions
separately for the two types of subnetworks
- Estimate the information exchange rate (IER/wIER) based on the dominant intrinsic
coupling modes

The novel results of our analysis include the following:

- Classification of the two groups based on the PD-comodulograms reached an accuracy of approximately 99%.
- Using linear regression analysis, we detected four cross-frequency pairs with δ as the dominant phase modulator and (δ, θ), (δ, β), (δ, γ_1_), (δ, γ_2_) as the most significant features that can classify the two groups correctly.

Considering that typical structural imaging alone might fail to indicate the development of mTBI, Huang and co-workers (Huang et al. 2009) proposed integrating MEG/MRI scans with DTI. They demonstrated the superiority of the bimodal approach over MRI and DTI alone in efficiently detecting mTBI by correlating MEG slow waves with fractional anisotropy in DTI and, thus, linking functional disturbances with specific cortical grey-matter areas. Interestingly, they also found that in some abnormal MEG recordings, δ waves were not accompanied by changes in fractional anisotropy, indicating the superiority of MEG in detecting mTBI, even in the absence of structural changes (Huang et al., 2009). Recently, we observed δ hyperactivity in right frontal brain areas in mTBI subjects using a novel complexity index to analyze MEG activity (Antonakakis et al., 2016b).

Many other analysis procedures applied to both EEG (e.g., Arakaki et al., 2017) and MEG (e.g., Zouridakis et al., 2017) recordings have reported altered functional connectivity in mTBI, which, in some cases, was correlated with the severity of the disease (Castellanos et al., 2010; for a review see Talavage et al., 2016). Adopting Granger causality as a connectivity measure, Zouridakis and co-workers (Zouridakis et al., 2012) reported that brain networks of mTBI patients exhibited fewer long-range connections compared to healthy controls, a few weeks after mTBI. Two more recent studies demonstrated the sensitivity of resting-state MEG recordings to detect abnormal connectivity in TBI (Tarapore et al., 2013) and mTBI (Da Costa et al.,2014), while a strong correlation between structural and functional features has been revealed between δ waves (MEG) and axonal injury (DTI) (Huang et al., 2009, 2014; Mvula et al., 2017). Along similar lines, recent studies have detected hyper-synchronization in mTBI subjects in the δ band (Dunkley et al., 2015; Li et al., 2015). Furthermore, Li et al. (2015) demonstrated an over-activation of intracranial sources in mTBI in δ, θ, and low α frequency bands compared to controls. Analysis of evoked potentials and ongoing MEG activity obtained from mTBI patients and controls across three repeat sessions scheduled approximately two and four weeks apart from the initial session showed that working memory processing in mTBI subjects does improve overtime (Arakaki et al., 2017); however, functional brain connectivity patterns do not recover at the rate that we might have expected (Zouridakis et al., 2016).

In two of our recent studies, we demonstrated effective discrimination of mTBI patients from controls by combining brain networks and machine learning techniques using phase-locking estimators (Dimitriadis et al., 2015a). In a follow-up analysis focusing on inter-frequency coupling (Antonakakis et al., 2016a), mTBI demonstrated lower integration and weaker local and distant connections compared to controls (see also a review by Rapp et al., 2015). Furthermore, in a dynamic fashion of the inter-frequency coupling, mTBI showed higher segregation and slower micro state transitions and complexity compared to controls (Antonakakis et al, 2016b, c).

Many recent neuroimaging studies have suggested that both structural and functional brain connectivity networks exhibit “small-world” characteristics, whereas recent studies based on structural DTI data have also revealed a “rich-club” organization of brain networks. Rich-club hubs of high-connection density tend to connect more often among themselves compared to nodes of lower density (van den Heuvel et al., 2013). In a recent study, we adopted an attack strategy to deduce the dominant type of network model (in terms of RC or SW organization) that best describes MEG resting-state networks for control and mTBI subjects (Antonakakis et al., 2015). RC nodes play a significant role in the information flow among anatomically distant brain subsystems that oscillate in a prominent type of interaction. For that reason, cross-frequency coupling plays a key role on the integration of the brain functionality (Antonakakis et al., 2016a; Dimitriadis et al., 2015b, c, 2016a, b). Thus, it seems necessary to focus on how this integration is affected by various brain diseases and disorders, taking into account the dominant types of interactions in the brain (Dimitriadis et al., 2015c, 2016b) but also the subdivisions of the functional brain network based on its RC organization (van den Heuvel and Sporns, 2011; Mišić et al., 2014; Bullmore and Sporns, 2012).

In our previous studies, we estimated intra-frequency functional brain networks for both mTBI and control groups (Dimitriadis et al., 2015a), while for the first time crossfrequency interactions were explored in a more recent mTBI study (Antonakakis et al., 2016a). Here, we detected the dominant type of interaction for each pair of MEG sensors and then the functional brain network was divided into two subsystems, namely, RC hubs and non-RC hubs. We then estimated the ratio of probability distribution of dominant intrinsic coupling modes within the RC hub subnetwork and between the RC hubs and the rest of the network. This stratification of the functional brain networks with the incorporation of dominant intrinsic coupling modes succeeded to discriminate mTBI from healthy controls with considerable success. Comparing with related recent results, Dimitriadis et al., (2015a) examined the metric of relative power but with low percentage of mTBI detection, while Antonakakis et al., (2016b) achieved more accurate classification results at the level of 97.5% using the complexity index and statistical differences in the same network areas (Fig. 1). The present study attempts to go beyond classification, towards a neuro-functional modeling of mTBI effects. Additionally, by adopting a general linear regression model of binomial distribution, we showed that using as independent variables the ratio of four frequency pairs, with δ as the phase modulator, i.e., (δ, θ), (δ, β), (δ, γ_1_), and (δ, γ_2_), could accurately predict the subject group (dependent variable). This demonstrated a hyperactivity and an increased rate of information exchange within the RC hub in mTBI subjects, which can be interpreted as a compensatory mechanism of the injury. It seems that this hyperactivity within the RC subnetwork is δ-phase mediated and can be interpreted as a mechanism for balancing of loss of connections and the reduction in the theoretical exchange of information can be expressed through IER and wIER metrics in the mTBI subjects.

The current sensor-level MEG analysis provided significant results and, thus, a future analysis at the source level is necessary to confirm the current findings, possibly by combining structural and functional data to estimate brain source connectivity (MartÍn-Buro et al, 2016). We expect that because of the negligible effects of head conductivity (Hämäläinen et al., 1993) on MEG recordings, the outcomes might be similar. An important improvement of our analysis would be the adoption of a dynamic functional connectivity scheme (Dimitriadis et al., 2009, 2010a, 2012a, b, c, 2013b, 2015c; Pang et al., 2016) summarized in functional connectivity microstates (Dimitriadis et al., 2013b, c, 2015b, d) and network microstates (Dimitriadis et al., 2013b, c, 2015b; Antonakakis et al., 2016c) which can provide more accurate results on a millisecond basis. Moreover, we plan to study the repertoire and the temporal variability of dominant intrinsic coupling modes in the two subject groups to further understand the effects of mTBI via graph theory at resting state (Dimitriadis et al., 2016b). Along this direction, we are attempting to link functional with structural networks and features form fractional anisotropy to behavioural data (Mvula et al., 2017), as well as to access sensitivity of functional states and their coupling to the recovery period (Tarapore et al., 2013; Arakaki et al., 2017).

## Acknowledgement

The current study is part of a larger project, the Integrated Clinical Protocol, conducted by the Investigators and staff of The Mission Connect Mild Traumatic Brain Injury Translational Research Consortium and supported by the Department of Defense Congressionally Directed Medical Research Program W81XWH-08-2-0135. SID was supported by MRC grant MR/K004360/1 (Behavioural and Neurophysiological Effects of Schizophrenia Risk Genes: A Multi-locus, Pathway Based Approach) and by EU-UK COFUND FELLOWSHIP.

The authors want to dedicate current study in remembrance to Prof. Sifis Micheloyannis, who is no longer with us, for his important contribution in neuroscience. A renown scientist with passion for research and eagerness for helping young scientists.

## Supplementary Material

### 1. Patient demographics

The current study is part of a larger mTBI project (Levin, 2009) supported by the Department of Defense (DoD). The subjects included in this analysis included a group of 30 right-handed patients with mTBI (29.33 ± 9.2 years of age) from the DoD project and a group of 50 age-matched neurologically intact controls (29.25 ± 9.1 years of age) drawn from a database that was being assembled as a normative data repository at UTHSC-Houston. The definition of mTBI used followed the guidelines of DoD (Assistant Secretary, 2007) and the American Congress of Rehabilitation Medicine (Kay et al., 1993). Mild TBI subjects were recruited from the Emergency Departments (EDs) of two Level 1 trauma centers and one Level III community hospital in a large ethnically diverse southwestern metropolitan area. Subjects were recruited by healthcare professionals (RN, MD, EMT-P) who had clinical experience with brain injury patients, knowledge of research, and excellent interpersonal and problem-solving skills. Screening occurred through review of data in the EDs electronic healthcare system (EHS), consultation with ED staff, and subject interviews. Special permission was obtained from the institutional IRBs to administer the Galveston Orientation and Amnesia Test (GOAT) (Levin et al., 2008) prior to obtaining informed consent to identify cognitive impairment that would preclude provision of informed consent. All subjects showed GOAT scores of 75 or greater and so have provided informed consent.

Inclusion criteria for the mTBI subjects included age 18-50 years, injury occurring within the preceding 24 hours, presence of a head injury (documented in medical records and/or verified by witnesses), Glasgow Coma Scale (GCS) (Teasdale & Jennett, 1974) score 13-15, loss of consciousness <30 minutes including 0 minutes, post-traumatic amnesia <24 hours including 0 minutes, and a negative head computed tomography (CT) scan. Exclusion criteria included a score on the Abbreviated Injury Scale (AIS) >3 for any body part, history of significant pre-existing disease (e.g., psychotic disorder, bipolar disorder, post-traumatic stress disorder (PTSD) diagnosed by a psychiatrist or psychologist, past treatment for alcohol dependence or substance abuse), blood alcohol level >80 mg/dL at the time of consent, documentation of intoxication, left-handed ness, and contraindications for MRI (including claustrophobia and pregnancy). Previous head injury requiring hospitalization or ED treatment was also an exclusion criterion. The demographics of mTBI subjects and the location of injury are given in Table S3.

The normative data repository included neurologically intact right-handed adults recruited from the University of Texas Medical School (UTMS) population (medical students and fellows). Handedness was assessed using the Edinburgh Handedness Inventory (Oldfield, 1971). Participants were screened, using self-report, for medication affecting the neurophysiological activity of the brain, as well as metallic implants, such as dental crowns, which affect the MEG evoked fields. Previous head injury, history of neurological or psychiatric disorder, substance abuse, and extensive dental work and implants incompatible with MEG were exclusion criteria for the control subjects. The project was approved by the Institutional Review Boards at the participating institutions and the Human Research Protection Officials review of research protocols for DoD. All procedures were compliant with the Health Insurance Portability and Accountability Act (HIPAA).

**Table S3.**
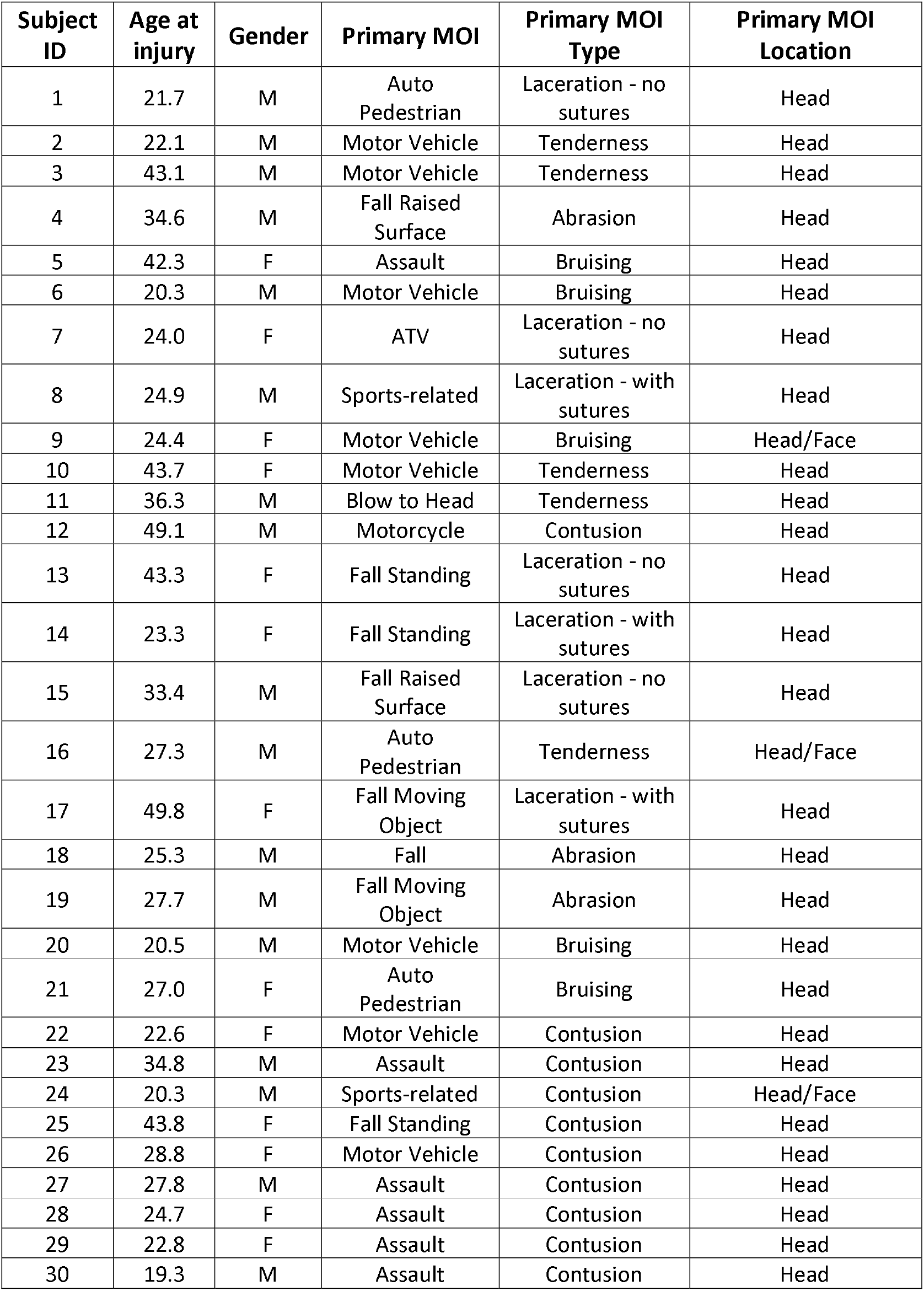
Subject demographics, location, and mode of impact (MOI) for the mTBI group.

| Subject ID | Age at injury | Gender | Primary MOI | Primary MOI Type | Primary MOI Location |
| --- | --- | --- | --- | --- | --- |
| 1 | 21.7 | M | Auto Pedestrian | Laceration - no sutures | Head |
| 2 | 22.1 | M | Motor Vehicle | Tenderness | Head |
| 3 | 43.1 | M | Motor Vehicle | Tenderness | Head |
| 4 | 34.6 | M | Fall Raised Surface | Abrasion | Head |
| 5 | 42.3 | F | Assault | Bruising | Head |
| 6 | 20.3 | M | Motor Vehicle | Bruising | Head |
| 7 | 24.0 | F | ATV | Laceration - no sutures | Head |
| 8 | 24.9 | M | Sports-related | Laceration - with sutures | Head |
| 9 | 24.4 | F | Motor Vehicle | Bruising | Head/Face |
| 10 | 43.7 | F | Motor Vehicle | Tenderness | Head |
| 11 | 36.3 | M | Blow to Head | Tenderness | Head |
| 12 | 49.1 | M | Motorcycle | Contusion | Head |
| 13 | 43.3 | F | Fall Standing | Laceration - no sutures | Head |
| 14 | 23.3 | F | Fall Standing | Laceration - with sutures | Head |
| 15 | 33.4 | M | Fall Raised Surface | Laceration - no sutures | Head |
| 16 | 27.3 | M | Auto Pedestrian | Tenderness | Head/Face |
| 17 | 49.8 | F | Fall Moving Object | Laceration - with sutures | Head |
| 18 | 25.3 | M | Fall | Abrasion | Head |
| 19 | 27.7 | M | Fall Moving Object | Abrasion | Head |
| 20 | 20.5 | M | Motor Vehicle | Bruising | Head |
| 21 | 27.0 | F | Auto Pedestrian | Bruising | Head |
| 22 | 22.6 | F | Motor Vehicle | Contusion | Head |
| 23 | 34.8 | M | Assault | Contusion | Head |
| 24 | 20.3 | M | Sports-related | Contusion | Head/Face |
| 25 | 43.8 | F | Fall Standing | Contusion | Head |
| 26 | 28.8 | F | Motor Vehicle | Contusion | Head |
| 27 | 27.8 | M | Assault | Contusion | Head |
| 28 | 24.7 | F | Assault | Contusion | Head |
| 29 | 22.8 | F | Assault | Contusion | Head |
| 30 | 19.3 | M | Assault | Contusion | Head |

### 2. Phase to Amplitude Coupling

Given a multidimensional array of time series **X**, PAC was calculated for the data from each sensor X_i_ and between pairs of sensors X_i_ and X_j_, with i, j = 1 … 248, using mutual information (MI) (Tsiaras et al., 2011; Bullmore et al., 2011). First, we extracted the low-frequency phase (f_l,i_) of the i-th component φ*_f_l_, i_* and the high-frequency amplitude (f_h,j_) of the j-th component A_fh,j_ using the Hilbert transformation (HT) (Claerbout, 1985). More specifically, f_h_ covered frequencies from θ to γ_2_, whereas f_l_ varied from δ to γ_1_. The cutoff frequency of the lowpass filter was higher than the cutoff of the highpass, so that the two filtering operators preserved a common bandpass interval. Since the power spectrum of A_fh,j_ preserved only a small portion of the very high frequencies, we bandpass filtered it to match the frequency range of φ*_f_l,i__*. Then, the phase of A_fh,j_, denoted by 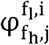, was extracted by a second HT. Finally, the estimation of PAC_*fc*_ was performed through Eq. (1), where 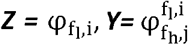, and *f_c_* = (*f_l_,f_h_*) = [(δ,θ), …, (γ_1_, γ_2_)]

To compute the Cross Frequency Functional Connectivity Graphs (CFCG) - PAC values, we used the HT to estimate the phase (φ*_f,i_*) and amplitude (*A_f,i_*) of every *X_f,i_*, separately in each frequency band using

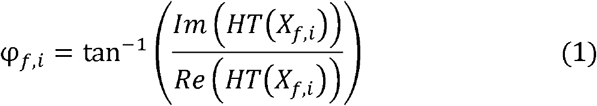

and

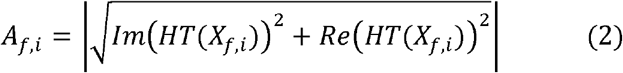

where Im(HT(X_f,i_)) and Re(HT(X_f,i_)) are the imaginary and real parts of HT(X_f,i_), respectively. We then applied a band-pass filter to A_f,i_ using the same filter parameters used to extract X_f1,i_, which resulted in a new time series, A_fh,f1,i_. A second HT was then used to extract the phases of the f_1_filtered f_h(high)_ amplitude envelope (φ_fh,f1,i_) (Voytek et al., 2010). The estimation of PAC between the phase of low frequency f_1_,φ_f_1_,i_ and the amplitude of the high frequency f_h_,φ_fh,f_1_,i_ between two sensors X_i_ and X_j_, is given by Eq. (1) in the main text, where Z = φ_f_1_,i_ and Y = φ_fh,f_1_,i_.

## Footnotes

1 http://users.auth.qr/~stdimitr/software.html

2 https://sites.google.com/site/bctnet/

## References

Alawieh A, Sabra Z, Sabra M, Tomlinson S, & Zaraket FA, (2015) A Rich-Club Organization in Brain Ischemia Protein Interaction. NetworkSci Rep 5:13513.

Antonakakis M, Dimitriadis SI, Papanicolaou AC, Gouridakis G, Zervakis M (2016b), Improving the Detection of mTBI Via Complexity Analysis in Resting – State Magnetoencephalography. Conf Proc IEEE Im Sys Tech.

Antonakakis M, Dimitriadis SI, Zervakis M, Micheloyannis S, Rezaie R, Babajani-Feremi A, Gouridakis G, Papanicolaou AC (2016a), Altered cross-frequency coupling in resting-state MEG after mild traumatic brain injury. Int J Psychophysiol 102:1–11.

Antonakakis M, Dimitriadis SI, Zervakis M, Papanicolaou AC, Gouridakis G (2016c), Mining Cross-Frequency Coupling Microstates from Resting State MEG: An Application to Mild Traumatic Brain Injury. Conf Proc IEEE Eng Med Biol Soc.

Antonakakis M, Dimitriadis SI, Zervakis M, Rezaie R, Babajani-Feremi A, Micheloyannis S, Gouridakis G, Papanicolaou AC, (2015) Comparison of brain network models using cross-frequency coupling and attack strategies. Conf Proc IEEE Eng Med Biol Soc 7426–7429.

Arakaki X, Shoga M, Li L, Zouridakis G, Dawlaty J, Goldweber R, Harrington M, Alpha power during working memory is compromised in acute mild traumatic brain injury, 12th World Congress on Brain Injury, March 29-April 1, 2017, New Orleans, LA

Aru J, Priesemann V, Wibral M, Lana L, Pipa G, Singer W, and Vicente R (2015), Untangling cross-frequency coupling in neuroscience. Curr Opin Neurobiol 31:51–61.

Assistant Secretary DoD (2007), Traumatic Brain Injury: Definition and Reporting. Department of Defense.

Bassett DS, Bullmore ET, Meyer-Lindenberg A, Apud JA, Weinberger DR, & Coppola R (2009), Cognitive fitness of cost-efficient brain functional networks. Proc Natl Acad Sci, 106:11747–11752.

Benjamíni Y, & Hochberg Y (1995), Controlling the False Discovery Rate: A Practical and Powerful Approach to Multiple Testing. J R Statist Soc 57:289–300.

Bullmore E, & Sporns O (2012), The economy of brain network organization. Nat Rev Neurosci 13:336–349.

Canolty RT, Edwards E, Dalal SS, Soltani M, Nagarajan SS, Kirsch HE, Berger MS, Barbaro NM, Knight RT (2006), High gamma power is phase-locked to theta oscillations in human neocortex. Science 313(5793):1626–1628.

Castellanos NP, Paúl N, Ordóñez VE, Demuynck O, Bajo R, Campo P, Bilbao A, Ortiz T, del-Pozo F, Maest? F (2010), Reorganization of functional connectivity as a correlate of cognitive recovery in acquired brain injury. Brain 133:2365–2381.

Cortes C, Vapnik V, 1995. Support-vector networksMachLearn 20 (3), 273–297.

Crossley NA, Mechelli A, Ginestet C, Rubinov M, Bullmore ET, & McGuire P (2016), Altered Hub Functioning and Compensatory Activations in the Connectome: A Meta-Analysis of Functional Neuroimaging Studies in Schizophrenia. Schizophr Bull, 42:434–442.

Da Costa L, Robertson A, Bethune A, MacDonald MJ, Shek PN, Taylor MJ, & Pang EW (2015), Delayed and disorganised brain activation detected with magnetoencephalography after mild traumatic brain injury. J Neurol Neurosurg Psychiatry 86:1008–1015.

Delorme A, & Makeig S (2004), EEGLAB: an open source toolbox for analysis of single-trial EEG dynamics including independent component analysis. J Neurosci Methods 134:9–21.

Dimitriadis S, Sun Y, Laskaris N, Thakor N, Bezerianos A (2016a), Revealing cross-frequency causal interactions during a mental arithmetic task through symbolic transfer entropy: a novel vector-quantization approach. IEEE Trans Neural Syst Rehabil Eng 24(10): 1017–1028.

Dimitriadis SI, Laskaris NA, Bitzidou MP, Tarnanas I and Tsolaki MN (2015d), A novel biomarker of amnestic MCI based on dynamic cross-frequency coupling patterns during cognitive brain responses. Front Neurosci 9:350.

Dimitriadis SI, Laskaris NA, Del Rio-Portilla Y, Koudounis GC (2009), Characterizing dynamic functional connectivity across sleep stages from EEG. Brain Topography, 22:119–133.

Dimitriadis SI, Laskaris NA, Micheloyannis S (2015b), Dynamics of EEG-based Network Microstates unmask developmental and task differences during mental arithmetic and resting wakefulness. Cogn Neurodyn 9(4):371–387.

Dimitriadis SI, Laskaris NA, Simos PG, Fletcher JM, Papanicolaou AC (2016b), Greater repertoire and temporal variability of cross-frequency coupling (CFC) modes in resting-state neuromagnetic recordings among children with reading difficulties. Front Hum Neurosci 10:163.

Dimitriadis SI, Laskaris NA, Tzelepi A (2013b), On the quantization of time-varying phase synchrony patterns into distinct functional connectivity microstates (FCμstates) in a multi-trial visual ERP paradigm. Brain Topogr 26(3):397–409.

Dimitriadis SI, Laskaris, NA, Simos PG, Micheloyannis S, Fletcher JM, Rezaie R, Papanicolaou AC (2013a), Altered temporal correlations in resting-state connectivity fluctuations in children with reading difficulties detected via MEG. NeuroImage 83:307–31.

Dimitriadis SI, Sun Yu, Kwok K, Laskaris NA, Thakor N, Bezerianos A (2013c), A tensorial approach to access cognitive workload related to mental arithmetic from EEG functional connectivity estimates. Conf Proc IEEE Eng Med Biol Soc 2013:2940–3.

Dimitriadis SI, Sun Yu, Kwok K, Laskaris NA, Thakor N, Bezerianos A (2015c), Cognitive Workload Assessment Based on the Tensorial Treatment of EEG Estimates of Cross-Frequency Phase Interactions. Ann Biomed Eng 43(4):977–89.

Dimitriadis SI, Zouridakis G, Rezaie R, Babajani-Feremi A, Papanicolaou AC (2015a), Functional connectivity changes detected with magnetoencephalography after mild traumatic brain injury. NeuroImage: Clinical 9:519–531.

Dunkley BT, Da Costa L, Bethune A, Jetly R, Pang EW, Taylor MJ, & Doesburg SM (2015), Low-frequency connectivity is associated with mild traumatic brain injury. NeuroImage Clin 7:611–621.

Engel AK, Gerloff C, Hilgetag CC, Nolte G (2013) Intrinsic coupling modes: multiscale interactions in ongoing brain activity. Neuron 80(4):867–86.

Escudero J, Hornero R, Abásolo D, & Fernández A (2011), Quantitative evaluation of artifact removal in real magnetoencephalogram signals with blind source separation. Ann Biomed Eng 39:2274–2286.

Fornito A, Zalesky A, Pantelis C, & Bullmore ET (2012), Schizophrenia, neuroimaging and connectomics. NeuroImage, 62:2296–2314.

Hämäläinen M, Hari R, Ilmoniemi RJ, Knuutila J, Lounasmaa OV, (1993) Magnetoencephalography — theory, instrumentation, and applications to noninvasive studies of the working human brain. Rev Mod Phys 65:413–197.

He X, Cai D, Niyogi P (2005) Laplacian score for feature selection. Adv in Neural Inform Proc Systems.

Horn RA, Mathias R (1990), An analog of the Cauchy-Schwarz inequality for Hadamard products and unitarily invariant norms. SIAM J Matrix Anal & Appl 11(4):481–498

Huang MX, Nichols S, Baker DG, Robb A, Angeles A, Yurgil KA, Drake A, Levy M, Song T, McLay R, Theilmann RJ, Diwakar M, Risbrough VB, Ji Z, Huang CW, Chang DG, Harrington DL, Muzzatti L, Canive J, Christopher M, Edgar J, Chen YH, Lee RR (2014), Single-subject-based whole-brain MEG slow-wave imaging approach for detecting abnormality in patients with mild traumatic brain injury. NeuroImage Clin 5:109–119.

Huang MX, Theilmann RJ, Robb A, Angeles A, Nichols S, Drake A, D’Andrea J, Levy M, Holland M, Song T, Ge S, Hwang E, Yoo K, Cui L, Baker DG, Trauner D, Coimbra R, Lee RR (2009), Integrated imaging approach with MEG and DTI to detect mild traumatic brain injury in military and civilian patients. J Neurotrauma 26(8):1213–26.

Kay T (1993), Mild traumatic brain injury committee of the head injury interdisciplinary special interest group of the American Congress of Rehabilitation Medicine Definition of mild traumatic brain injury. J Head Trauma Rehabil 8:86–87.

Len TK, & Neary JP (2011), Cerebrovascular pathophysiology following mild traumatic brain injury. Clin Physiol Func Imaging, 31:85–93.

Levin HS (2009) Mission Connect Mild TBI Translational Research Consortium. Baylor College of Medicine Houston TX.

Levin HS, Mattis S, Ruff RM, Eisenberg HM, Marshall LF, Tabaddor K, High WM Jr, Frankowski, RF (1987), Neurαbehaviαral outcome following minor head injury: a three-center study. J Neurosurg, 66:234–243.

Li L, Mvula E, Arakaki X, Tran T, Harrington M, Zouridakis G Source Connectivity Analysis Can Assess Recovery of Acute Mild Traumatic Brain Injury Patients, 12th World Congress on Brain Injury, March 29 - April 1, 2017, New Orleans, LA

Li L, Pagnotta MF, Arakaki X, Tran T, Strickland D, Harrington M, Zouridakis G (2015), Brain activation profiles in mTBI: Evidence from combined resting-state EEG and MEG activity. Conf Proc IEEE Eng Med Biol Soc, 6963–6966.

Liu Y, Liang M, Zhou Y, He Y, Hao Y, Song M, Yu C, Liu H, Liu Z, Jiang T (2008), Disrupted small-world networks in schizophrenia. Brain 131:945–961.

MartÍn-Buro MC, Garcés P, Maest? F (2016) Test-retest reliability of resting-state magnetoencephalography power in sensor and source space. Hum Brain Mapp 37(1):179–90.

Micheloyannis S, Pachou E, Stam CJ, Breakspear M, Bitsios P, Vourkas M, Erimaki S, Zervakis M (2006), Small-world networks and disturbed functional connectivity in schizophrenia. Schizophr Res 87:60–66.

Mišić B, Sporns O, & McIntosh AR (2014), Communication efficiency and congestion of signal traffic in large-scale brain networks PLoS Comput Biol 10:el003427.

Mvula E, Li L, Arakaki X, Tran T, Harrington M, Zouridakis G Assessing Recovery of Acute Mild Traumatic Brain Injury Patients using Diffusion Tensor Imaging, 12th World Congress on Brain Injury, March 29 - April 1, 2017, New Orleans, LA

Oostenveld R, Fries P, Maris E, Schoffelen (2011), FieldTrip: Open Source Software for Advanced Analysis of MEG, EEG, and Invasive Electrophysiological Data. Comput Intell Neurosci, 2011:el56869.

Palva JM, & Palva S (2011), Roles of multiscale brain activity fluctuations in shaping the variability and dynamics of psychophysical performance. Prog Brain Res 193:335–350.

Rapp PE, Keyser DO, Albano A, Hernandez R, Gibson DB, Zambón RA, Hairston WD, Hughes JD, Krystal A, Nichols AS (2015), Traumatic Brain Injury Detection Using Electrophysiological Methods. Front Hum Neurosci 9:11.

Rubinov M, Sporns O (2010), Complex network measures of brain connectivity: uses and interpretations. NeuroImage, 52:1059–1069.

Senden M, Deco G, de Reus MA, Goebel R, & van den Heuvel MP (2014), Rich club organization supports a diverse set of functional network configurations NeuroImage, 96 174–182.

Stam CJ, Jones BF, Nolte G, Breakspear M, & Scheltens P (2007), Small-world networks and functional connectivity in Alzheimer’s disease Cereb Cortex 17:92–99.

Talavage TM, Nauman EA and Leverenz LJ (2016), The Role of Medical Imaging in the Recharacterization of Mild Traumatic Brain Injury Using Youth Sports as a Laboratory. Front Neurol 6:273.

Tarapore PE, Findlay AM, Lahue SC, Lee H, Honma SM, Mizuiri D, Mukherjee P (2013), Resting state magnetoencephalography functional connectivity in traumatic brain injury. J Neurosurg 118:1306–1316.

Theiler J, Eubank S, Longtin A, Galdrikian B, and Farmer J D (1992), Testing for nonlineaity in time series:the method of surrogate data. Physica D 85:77.

Tort ABL, Kramer MA, Thorn C, Gibson DJ, Kubota Y, Graybiel AM, & Kopell NJ (2008), Dynamic cross-frequency couplings of local field potential oscillations in rat striatum and hippocampus during performance of a T-maze task Proc Nat Acad Sci, 105:20517–20522.

Tsiaras V, Simos PG, Rezaie R, Sheth BR, Garyfallidis E, Castillo EM, & Papanicolaou AC (2011), Extracting biomarkers of autism from MEG resting-state functional connectivity networks. Comput Biol Med, 41:1166–1177.

Van de Heuvel MP, & Sporns O (2011), Rich-Club Organization of the Human Connectome. J Neurosci 31:15775–15786.

Van den Heuvel MP, Sporns O, Collin G, Scheewe T, Mandl RCW, Cahn W, Goñi J, Hulshoff Pol, HE, Kahn, RS (2013), Abnormal rich club organization and functional brain dynamics in schizophrenia. JAMA Psychiatry, 70:783–792.

Varela F, Lachaux JP, Rodriguez E, & Martinerie J (2001), The brainweb: Phase synchronization and large-scale integration. Nat Rev Neurosc 2:229–239.

Vértes, PE, & Bullmore ET (2015), Annual research review: Growth connectomics--the organization and reorganization of brain networks during normal and abnormal development. J Child Psychol Psychiatry, 56:299–320.

Voytek B, Canolty RT, Shestyuk A, Crone NE, Parvizi J, & Knight RT (2010), Shifts in Gamma Phase-Amplitude Coupling Frequency from Theta to Alpha over Posterior Cortex during Visual Tasks. Front Hum Neurosc 4:191.

Vrba J, & Robinson SE (2001), Signal processing in magnetoencephalography. Methods 25:249–271.

Watanabe T, & Rees G (2015), Age-associated changes in rich-club organization in autistic and neurotypical human brains. Sci Rep 5:16152.

Xu X, Zheng C, & Zhang T (2013), Reduction in LFP cross-frequency coupling between theta and gamma rhythms associated with impaired STP and LTP in a rat model of brain ischemia. Front Comput Neurosc 7:27.

Zouridakis G, Patidar U, Situ N, Rezaie R, Castillo EM, Levin HS, & Papanicolaou AC (2012), Functional connectivity changes in mild traumatic brain injury assessed using magnetoencephalography J Mech Med Biol 12:1240006.

Zouridakis G, Li L, Arakaki X, Tran T, Padhye N, and Harrington M Assessing Recovery of mTBI Patients using Functional Connectivity: A Resting State Magnetoencephalographic Study. 20th International Conference on Biomagnetism (BIOMAG2016), October 1–6, 2016, Seoul, Korea

## References

Assistant Secretary, D.o.D., 10–1-2007. Traumatic Brain Injury: Definition and Reporting. Department of Defense.

Bullmore ET, & Bassett DS (2011) Brain graphs: graphical models of the human brain connectome. Annu Rev Clin Psychol 7:113–140.

Claerbout, Jon F (1985). Fundamentals of Geophysical Data Processing with Applications to Petroleum Prospecting. Oxford, UK: Blackwell, 59–62.

Kay T (1993) Mild traumatic brain injury committee of the head injury interdisciplinary special interest group of the American Congress of Rehabilitation Medicine. Definition of mild traumatic brain injury. J Head Trauma Rehabil 8:86–87.

Levin HS (2009) Mission Connect Mild TBI Translational Research Consortium. Baylor College of Medicine Houston TX.

Levin HS, O’Donnell VM, & Grossman RG. (2008). The Galveston Orientation and Amnesia Test: A practical scale to assess cognition after head injury. J Nerv Ment Dis 167, 675–684.

Teasdale, G.M., Jennet, B.; 1974. Assessment of coma and impaired consciousness. Lancet 8184.

Teasdale, G.M., Jennet, B.; 1974. Assessment of coma and impaired consciousness. Lancet 8184.

Tsiaras V, Simos PG, Rezaie R, Sheth BR, Garyfallidis E, Castillo EM, & Papanicolaou AC (2011) Extracting biomarkers of autism from MEG resting-state functional connectivity networks. Comput Biol Med 41:1166–1177.

Voytek, B., Canolty, R. T., Shestyuk, A., Crone, N. E., Parvizi, J., & Knight, R. T. (2010). Shifts in Gamma Phase-Amplitude Coupling Frequency from Theta to Alpha Over Posterior Cortex During Visual Tasks. Frontiers in Human Neuroscience, 4. http://doi.org/10.3389/fnhum.2010.00191

